# A model to study NMDA receptors in early nervous system development

**DOI:** 10.1101/807115

**Authors:** Josiah Zoodsma, Kelvin Chan, David Golann, Ashwin Bhandiwad, Amalia Napoli, Guangmei Liu, Shoaib Syed, Harold Burgess, Howard Sirotkin, Lonnie P. Wollmuth

## Abstract

NMDA receptors (NMDARs) are glutamate-gated ion channels that play critical roles in neuronal development and nervous system function. Pharmacological antagonism is an invaluable tool to study NMDARs, but is experimentally limited. Here, we developed a model to study NMDARs in early development in zebrafish, by generating CRISPR-mediated lesions in the NMDAR genes, *grin1a* and *grin1b*, which encode the obligatory GluN1 subunits. While receptors containing *grin1a* or *grin1b* show high Ca^2+^ permeability, like their mammalian counterpart, *grin1a* is expressed earlier and more broadly in development than *grin1b*. Both *grin1a^−/−^* and *grin1b^−/−^* zebrafish are viable as adults. Unlike in rodents, where the *grin1* knockout is embryonic lethal, *grin1* double mutant fish (*grin1a^−/−^; grin1b^−/−^*), which lack all NMDAR-mediated synaptic transmission, survive until about 10 days post fertilization, providing a unique opportunity to explore NMDAR function during development and in generating behaviors. Many behavioral defects in the *grin1* double mutant larvae, including abnormal evoked responses to light and acoustic stimuli, prey capture deficits and a failure to habituate to acoustic stimuli, are replicated by short-term treatment with the NMDAR antagonist MK-801, suggesting they arise from acute effects of compromised NMDAR-mediated transmission. Other defects, however, such as periods of hyperactivity and alterations in place preference, are not phenocopied by MK-801, suggesting a developmental origin. Taken together, we have developed a unique model to study NMDARs in the developing vertebrate nervous system.

## INTRODUCTION

Nervous system function depends on the development of brain circuits that integrate appropriate cell types and establish proper connectivity. In the vertebrate nervous system, glutamate is the major excitatory neurotransmitter. At synapses that mediate rapid transmission, glutamate is converted into biological signals by ligand-gated or ionotropic glutamate receptors (iGluRs), including AMPA (AMPAR), kainate, and NMDA (NMDAR) receptor subtypes (Traynelis *et al.*, 2010). Because of its unique signaling properties, including high Ca^2+^ permeability, NMDARs are central to higher brain functions such as the plasticity underlining learning and memory (Hunt & Castillo, 2012; Paoletti *et al.*, 2013) and brain development (Cline & Haas, 2008; Gambrill & Barria, 2011; Chakraborty *et al.*, 2017). Due to the experimental limitations of pharmacological manipulation, especially in studies viewing long-term outcomes as occurs for many neurodevelopmental disorders, studying NMDARs in early circuit development is challenging. Adding to this challenge, the loss of NMDAR subunits critical to early brain development in murine models are embryonic lethal or die perinatally (Forrest *et al.*, 1994; Kutsuwada *et al.*, 1996; Sprengel *et al.*, 1998).

Zebrafish are a powerful model to study early nervous system development as they externally fertilize, which facilitates imaging of cell migration and circuit formation. Furthermore, zebrafish larvae exhibit complex behaviors that provide simple and robust readouts of higher nervous system function. Previous studies of NMDARs in zebrafish have generally used acute pharmacological manipulations, mainly MK-801, a general antagonist of NMDARs (Huettner & Bean, 1988; Traynelis *et al.*, 2010). These studies have highlighted the central role of NMDARs in complex brain functions and behaviors including associative learning (Sison & Gerlai, 2011), social interactions (Seibt *et al.*, 2011; Dreosti *et al.*, 2015), place preference (Swain *et al.*, 2004), and responses to acoustic stimuli and pre-pulse inhibition (Bergeron *et al.*, 2015). Although these studies are invaluable, they do not reveal how brain or circuit development impacts specific behaviors because acute antagonist applications only change brain function for a short developmental timepoint and can have off target effects.

NMDARs are heterotetramers composed of two obligate GluN1 subunits and typically two GluN2 (A-D) subunits (Paoletti *et al.*, 2013). In zebrafish, the obligatory GluN1 subunit is encoded by two paralogous genes: g*rin1a* and *grin1b*. The expression patterns of these genes have been described at early stages (Cox *et al.*, 2005); however, expression was assayed only during a limited time window, and any functional differences between paralogues are unknown. Here, we studied the developmental expression of *grin1a* and *grin1b* and the effect of their knockouts on early nervous system function. While their encoded proteins, GluN1a and GluN1b, show high Ca^2+^ permeability, like their mammalian counterparts, *grin1a* is expressed earlier in development and tends to be more widely expressed. Nevertheless, both *grin1a^−/−^* and *grin1b^−/−^* fish are viable as adults. Unlike rodents, *grin1* double mutant fish (*grin1a^−/−^*; *grin1b^−/−^*), which lack all NMDAR-mediated synaptic transmission, survive until about 10 days post fertilization (dpf), which is old enough to assess the impact of NMDAR deficits on behaviors. We took advantage of these fish to study the role of NMDARs in the development of numerous complex behaviors, including spontaneous and evoked movements, prey-capture, and habituation.

## MATERIALS AND METHODS

### Zebrafish maintenance and housing

Adult zebrafish strains were maintained at 28.5⁰C under a 13:11 hour light to dark cycle and were fed a diet of artemia and GEMMA micropellets. The wild-type strain used for all experiments was a hybrid wild-type background consisting of Tubingen long-fin crossed to Brian’s wild-type. The experiments and procedures were approved by the Stony Brook University IACUC.

### Whole-mount RNA in situ hybridization

The *grin1a* probe was previously described (Cox *et al.*, 2005). Probes for *grin1b* were generated by PCR amplification of 3 dpf wild-type cDNA using the following primers: *grin1b:* F (GAGtatttaggtgacactatagTCTGTGGACTAGCTGGCAAA) and R (GAGtaatacgactcactatagggATGGACGTTGCGTGTTTGTA)

Lower-case letters in forward and reverse primers indicate the T7 and SP6 RNA Polymerase binding sites, respectively. DIG-labeled anti-sense and sense probes were generated from PCR product.

Whole-mount RNA *in situ* hybridization was performed as described by Thisse et al. (Thisse *et al.*, 1993). Embryos were stained with nitro-blue tetrazolium and 5-bromo-4-chloro-3-indolyl phosphate (NBT/BCIP). Prior to photography, embryos were cleared with 2:1 benzyl benzoate to benzyl alcohol and then mounted in Canada Balsam with 2.5% methyl salicyclate. Embryos were imaged on a Zeiss Axioplan 2 microscope with Axiocam HRc mounted camera and AxioVision Software.

### CRISPR-Cas9 gene targeting and zebrafish microinjections

gRNA were designed through IDT custom gRNA Design Tool. gRNA were complexed to Cas9 protein, forming a Ribonucleoprotein (RNP), using the Alt-R^TM^ CRISPR-Cas9 System (IDT). 0.5 nL of RNP (25 pg of gRNA and 125 pg of Cas9 protein) were injected into the cell of pronased (5 mg/mL) 1-cell embryos. 10-20 embryos from each injection were assayed (by PCR) for lesions, and if detected, remaining embryos were grown into adulthood to be outcrossed and screened for germline transmission. All mutations were outcrossed at least twice to minimize the effect of off-target endonuclease activity on behavioral phenotypes.

### Zebrafish genotyping

Larval zebrafish viability was assayed at 6 dpf. Adult zebrafish viability and size were assayed after 2 months post-fertilization. The following primers were used to screen the CRISPR mutations by PCR and for subsequent genotyping:

*grin1a:* F (ATTAGGAATGGTGTGGGCTGGC) and R (GGTGATGCGCTCCTCAGGCC)

*grin1b:* F (GGTGCCCCTCGGAGCTTTTC) and R (GGAAGGCTGCCAAATTGGCAGT)

### Reverse transcript PCR (rtPCR)

RNA were extracted from pools of 4 larvae at 3 dpf from homozygous mutant or homozygous wildtype intercrosses using Trizol (Invitrogen). cDNA was synthesized from 0.2 ng of RNA using Superscript II Reverse Transcriptase (Invitrogen). The following primers were used to screen the cDNA by PCR:

*grin1a:* F (ATTAGGAATGGTGTGGGCTGGC) and R (ATGAATTTGTCTGATGGGTTCCT)

*grin1b:* F (GTGCCCCTCGGAGCTTTTCG) and R (AGGAAGGCTGCCAAATTGGCA)

### Whole-cell electrophysiology

Human embryonic kidney 293 (HEK293) cells were grown in Dulbecco’s modified Eagle’s medium (DMEM), supplemented with 10% FBS, for 24 h before transfection. Zebrafish cDNA constructs were synthesized from GenScript in a pcDNA3.1(+)-p2A-eGFP vector. Zebrafish (zGluN1a/b) and rat (rGluN2A) NMDAR-encoding cDNA constructs, were co-transfected into HEK293 cells along with a separate peGFP-Cl construct at a ratio of 4:4:1 (N1:N2:eGFP) for whole-cell or macroscopic recordings using X-tremeGene HP (Roche, 06-366). HEK293 cells were bathed in medium containing the GluN2 competitive antagonist DL-2-amino-5-phosphopentanoic acid (APV, 100 µM, Tocris) and Mg^2+^ (100 µM). All experiments were performed 24-48 h post-transfection.

Whole-cell currents were recorded at room temperature (20–23°C) using an EPC-10 amplifier with PatchMaster software (HEKA), digitized at 10 kHz and low-pass filtered at 2.9 kHz (−3 dB) using an 8 pole low pass Bessel filter 4. Patch microelectrodes were filled with an intracellular solution (mM): 140 KCl, 10 HEPES, 1 BAPTA, 4 Mg^2+^-ATP, 0.3 Na^+^-GTP, pH 7.3 (KOH), 297 mOsm (sucrose). Our standard extracellular solution consisted of (mM): 150 NaCl, 2.5 KCl, and 10 HEPES, pH 7.2 (NaOH). Currents were measured within 10 min of going whole cell.

External solutions were applied using a piezo-driven double barrel application system. For agonist application, one barrel contained the external solution +0.1 mM glycine, whereas the other barrel contained both 0.1 mM glycine and 1 mM glutamate. For display, NMDAR currents were digitally refiltered at 500 Hz and resampled at 1 kHz.

### Pharmacology

MK-801 (20 mM) was diluted to in DMSO. For all experiments, unless otherwise noted, final concentration used was 20 μM in 0.1% DMSO.

### Quantitative PCR (qPCR)

RNA was extracted as stated above. Pools of 4 larvae were used at 3 dpf, while pools of 2 larvae were used at 5 dpf. qPCR was carried out on a LightCycler 480 (Roche) using PerfeCTa ® SYBR ® GreenFastMix ® (QuantaBio). Total RNA from each sample was normalized to β-actn. In each experiment, 3 pools of cDNA from 2-4 embryos were run in duplicate for each genotype. The following primers were used to screen the cDNA by PCR (Menezes 2015): *grin1a:* F (ATAAAGACGCCCGCAGGAAGC) and R (CGTGCTGACAGACGGGTCCGAC) *grin1b:* F (AATGCAGCTGGCCTTTGCAGC) and R (CTCTTGATGTTGGAGGCCAGGTTG)

### Live imaging

Zebrafish larvae were immersed in 3% Methylcellulose, after anesthetization in ice water, and images were taken on a Zeiss Discovery V.20. *grin1* double mutant numbers were enhanced through use of a behavior screen at 6 dpf.

### Spontaneous and evoked movement paradigms

Behavioral assays were performed on multiple clutches from different parents to minimize the effects of genetic background. Embryos were kept in 150 mm Petri dishes in egg water (6 g of synthetic sea salt, 20 mL of methylene blue (1 g/L) solution in to 20 L of water pH 7) at concentrations under 100 larvae per plate.

#### Locomotive behavior

At 48 hpf, embryos were transitioned to 24-well behavior plates, 1 embryo per well in 1.5 mL of egg water. Behavior was recorded between 4 and 6 dpf using a Zebrabox imaging system (Viewpoint Life Sciences, France) and tracked with automated video-tracking software (Zebralab; Viewpoint Life Sciences, France).

### Prey-capture assay and Paramecium maintenance

*Paramecium multimicronucleatum* was obtained from Carolina Biological Supply Company and were maintained at room temperature with new cultures generated every 3 weeks. Paramecia from cultures of 2-5 weeks of age were filtered and moved to a 35 mm Petri dish on the day of the assay.

#### Prey-capture assay

Zebrafish embryos were kept in 150 mm Petri dishes in egg water at concentrations under 100 per plate. Individual larvae were assayed at 7 dpf. 20-60 paramecia were added to individual 35 mm Petri dishes filled with 3 mL of either system or egg water. 50 frames (~2.5 seconds) of paramecia movement were recorded using an Excelis MPX5C-PRO camera with version Mosaic V2.0 software at the beginning and end of each trial, which lasted 1.5 hours. These videos were processed using ImageJ (Gahtan *et al.*, 2005). Paramecia-trace images were manually counted by a blind viewer. Multiple control plates containing only paramecium were used in each assay to account for any decrease in paramecia viability. Results were expressed as a ratio of the proportion of paramecia eaten normalized to their respective control. Preliminary assays showed decreased paramecia survival in the presence of MK-801 (data not shown). To ensure paramecia survival, we exposed wild type larvae to either MK-801 (20 μM in 0.1% DMSO) or control (0.1% DMSO) for 2 hours prior to the start of the feeding assay, after which the larvae were moved to untreated system water for the trial.

### Habituation

#### Behavioral analysis

Individual fish were placed in 9-well ~ 1 cm^2^ gridded plates and tested for changes in short latency startle responses (SLC) to acoustic/vibrational stimuli and habituation of SLC. Pulsed acoustic stimuli were delivered using a Type 3810 minishaker (Bruel-Kjaer) and controlled using a BNC-2110 DAQ board (National Instruments). Behavioral responses were imaged at 1000 frames/sec using a DRS Lightning high speed camera (DEL Imaging) and analyzed using FLOTE software (Burgess & Granato, 2007). SLC responses were assigned based on response latency and kinematics. Control larvae were wildtype and heterozygous siblings from the same clutch as mutants. We assigned larvae to each group by genotyping after behavioral analysis.

For SLC experiments, each plate of fish was presented with 20 presentations each of (1) high intensity startle stimuli (37 ± 1 dB re. m/s^2^) and a (2) low intensity startle stimuli (19 ± 1 dB re. m/s^2^). These 80 stimuli were delivered in pseudorandom order with a 30 s interval between consecutive stimulus presentations.

For habituation experiments, larvae were presented with high intensity startle stimuli with 1 s interval between stimulus presentations. This paradigm has demonstrated rapid habituation, with SLC decreases of up to 80% after 20 stimulus presentations (Wolman *et al.*, 2011). Due to the stochasticity of individual-level data, we analyzed responsiveness in 4 epochs of 10 stimuli. Habituation was quantified as the percent decrease in SLC responsiveness in the last epoch relative to the first epoch.

### Statistics

Data analysis was performed using IgorPro, Excel and MiniTab 18. All average values are presented as mean ± SEM. The number of replicates is indicated in the figure legends. In instances where we were only interested in whether outcomes were statistically different from wild type or an appropriate control, we used an unpaired two-tailed Student’s t-test to test for significance. In instances where we were interested in how multiple groups varied from each other, we used a one-factor ANOVA and followed with a post-hoc Tukey’s test. Statistical significance was set at *p* < 0.05.

## RESULTS

Functional NMDARs are heterotetramers composed of two obligatory GluN1 subunits and typically some combination of GluN2 (A-D) subunits. Initially, we characterized the development expression of the zebrafish *grin1* (GluN1) paralogues, *grin1a* and *grin1b*.

### Differential expression of zebrafish *GRIN1* paralogues, *grin1a* and *grin1b*, in early development

*grin1a* is expressed in the brain, retina and spinal cord at 1 day post-fertilization (dpf) with expression becoming more robust by 2 dpf (Cox *et al.*, 2005). Over the same period, *grin1b* expression is much weaker (Cox *et al.*, 2005). To further define their expression, we looked at *grin1a* and *grin1b* expression using RNA *in situ* hybridization over a wider developmental range (1, 3 & 5 dpf) (Figure 1). For these experiments, we used the same *grin1a* probe as used previously (Cox *et al.*, 2005), but redesigned the *grin1b* probe to reduce background (see Material & Methods).

**Figure 1.**
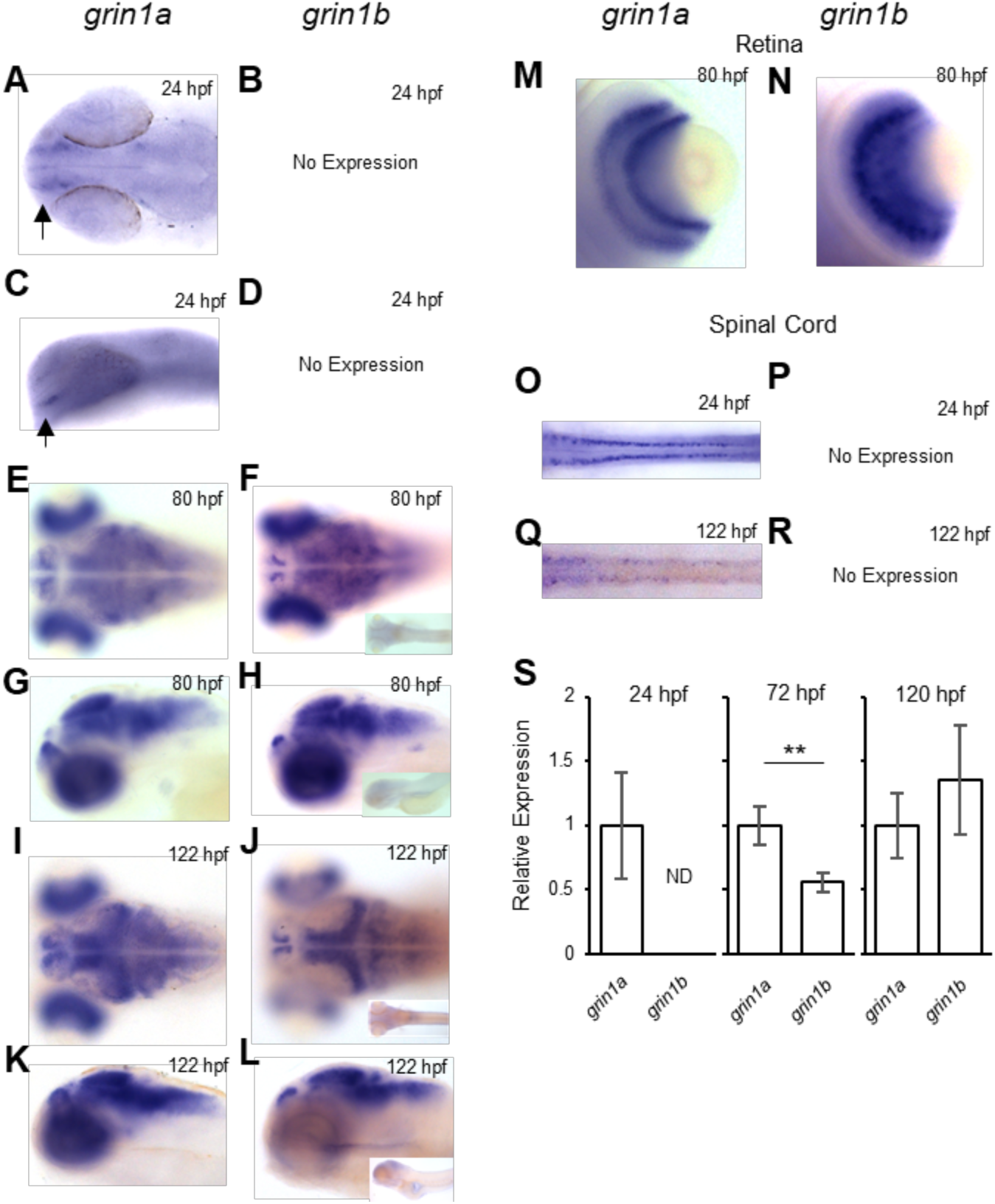
Expression of *grin1a* and *grin1b* in the zebrafish nervous system. Whole mount *in situ* hybridization of *grin1a* and *grin1b* at 24 hpf **(A-D)**, 80 hpf **(E-H)** and 122 hpf **(I-L)** showing dorsal (**A, E, F, I, J**) and lateral (**C, G, H, K, L**) views. Insets in **(F), (H), (J)**, and **(L)** are sense probes of *grin1b*. Arrows in **(A)** and **(C)** indicate limited bilateral expression. **(M-N)** Dorsal view of the retina at 80 hpf for *grin1a* **(M)** and *grin1b* **(N)**. **(O-R)** Dorsal view of the spinal cord at 24 hpf **(O & P)** and 120 hpf **(Q & R)** for *grin1a* and *grin1b*. **(S)** Relative rt-qPCR expression levels of *grin1a* and *grin1b* at 24 (n = 9), 72 (n = 9) and 120 (n = 6) hpf. ND = not detected. Results are normalized to *grin1a* expression at each individual timepoint. *\*\*p < 0.01, ratio paired t-test*.

Consistent with earlier results, we observed expression of *grin1a* at 24 hours post fertilization (hpf) in the telencephalon (Figures 1A & 1C, arrows) and in the spinal cord (Figure 1O). We could not detect any *grin1b* transcripts at this developmental stage (Figures 1B & 1D). By 80 hpf, *grin1a* and *grin1b* are expressed more broadly throughout the brain and in distinct domains in the retina (*grin1a*: Figures 1E, 1G, & 1M; *grin1b*: Figures 1F, 1H, & 1N). At 122 hpf, *grin1a* and *grin1b* are expressed in the telencephalon, posterior optic tectum, hindbrain (*grin1a*: Figures 1I & 1K; *grin1b*, Figures 1J & 1L) and in the retina (data not shown).

Although both *grin1a* and *grin1b* have overlapping expression, there are notable differences. In the spinal cord, *grin1a* is robustly expressed between 24 and 122 hpf, but we could not detect any *grin1b* expression at these stages (Figures 1O-1R). In the telencephalon and optic tectum, grin1a is broadly expressed at 122 hpf, whereas *grin1b* has a more discrete expression pattern (*grin1a*: Figures 1I & 1K; *grin1b*, Figures 1J & 1L).

The RNA *in situ* hybridization experiments suggest that *grin1a* is expressed at higher levels than *grin1b*. To directly test this idea, we carried out qPCR (Figure 1S) to measure levels of both transcripts. As expected, *grin1a* was readily detected at 24 hpf, whereas *grin1b* could not be detected (Figure 1S, left panel). In addition, at 72 hpf, the expression of *grin1a* was significantly greater than that of *grin1b* (Figures 1S, middle panel), but by 120 hpf, we could not detect any expression differences (Figures 1S, right panel). In summary, *grin1a* is expressed earlier than *grin1b* and shows a more robust expression pattern through 3 dpf.

### NMDARs containing zebrafish GluN1 paralogues are highly Ca^2+^ permeable like their mammalian counterparts

NMDARs are ligand-gated ion channels (Paoletti *et al.*, 2013). To define potential functional differences between zebrafish paralogues, we expressed *grin1a* and *grin1b* in a heterologous expression system (see Materials & Methods). To reduce potential variations with the GluN2 subunit, we expressed zebrafish GluN1a (zGluN1a) or GluN1b (zGluN1b) with rat GluN2A (rGluN2A) (Figure 2). As a reference, we compared these recordings to those for rat GluN1 (rGluN1) coexpressed with rGluN2A (rGluN1/rGluN2A).

**Figure 2.**
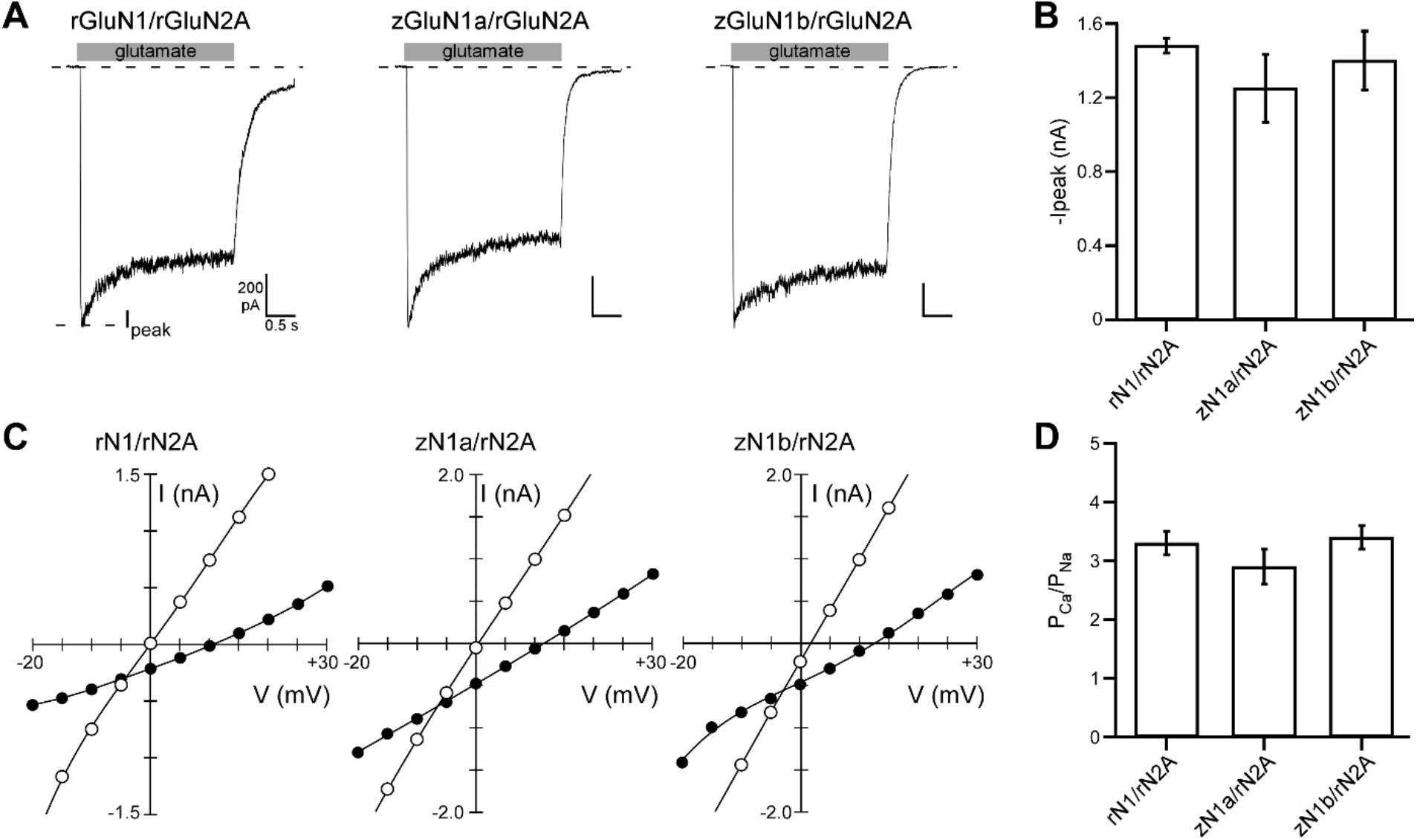
Zebrafish paralogues show functional currents and high Ca^2+^ permeability. **(A)** Whole-cell glutamate-activated currents recorded in HEK 293 cells in response to sustained (2.5 sec) glutamate applications (1 mM, gray bar) to measure NMDAR currents for various GluN1 constructs: rat GluN1 (rGluN1) (*left panel*); and zebrafish GluN1a (zGluN1a) (*middle panel*) or GluN1b (zGluN1b) (*right panel*). All GluN1 constructs were co-expressed with rat GluN2A (rGluN2A). Peak current amplitudes (I_peak_) were measured at the beginning of the glutamate application. Holding potential, −70 mV. Scale bar values are indicated in left panel. **(B)** Bar graph (mean ± SEM) of peak current amplitudes (I_peak_) for the various constructs (rGluN1, n = 10; zGluN1a, n = 6; zGluN1b, n = 7). None of the values were significantly different. **(C)** Measuring Ca^2+^ permeability. Current-voltage (IV) relationships in an external solution containing high Na^+^ (140 mM) and 0 added Ca^2+^ (open circles) or 10 mM Ca^2+^ (solid circles). The 0 Ca^2+^ IV is the average of that recorded before and after the 10 mM Ca^2+^ recording. We used changes in reversal potentials (□E_rev_) to calculate P_Ca_/P_Na_ (Jatzke *et al.*, 2002). **(D)** Relative calcium permeability (P_Ca_/P_Na_) (mean ± SEM) calculated from □E_rev_s for rat rGluN1/rGluN2A (8.3 ± 0.4 mV, n = 8), zGluN1a/rGluN2A (7.9 ± 0.7 mV, n = 9), or zGluN1b/rGluN2A (8.6 ± 0.5 mV, n = 8). None of the values were significantly different.

rGluN1/rGluN2A shows strong expression in HEK293 cells (Amin *et al.*, 2017) (Figure 2A, left panel). NMDARs containing zGluN1a (Figure 2A, center panel) or zGluN1b (Figure 2A, right panel) also showed robust current amplitudes that were indistinguishable from rGluN1 (Figure 2B).

A critical physiological property of NMDARs is their high Ca^2+^ permeability (Paoletti *et al.*, 2013), which is dominated by GluN1 (Burnashev *et al.*, 1992; Watanabe *et al.*, 2002). We therefore assayed the Ca^2+^ permeability (P_Ca_), relative to Na^+^ (P_Ca_/P_Na_), of NMDARs containing zGluN1a or zGluN1b (Figures 2C & 2D). To measure P_Ca_/P_Na_, we measured changes in reversal potentials going from a high Na^+^ solution without Ca^2+^ to the same solution containing 10 mM Ca^2+^ (see Materials & Methods)(Jatzke *et al.*, 2002; Amin *et al.*, 2018). NMDARs containing zGluN1a or zGluN1b showed a high Ca^2+^ permeability that is indistinguishable from that of NMDARs containing rGluN1 (Figure 2D). Thus, zebrafish NMDARs, regardless of the GluN1 paralogue, are highly Ca^2+^ permeable, like their mammalian counterpart (Wollmuth, 2018).

### Generation of lesions in *grin1a* and *grin1b*

To study the requirements for NMDAR-mediated transmission in the developing zebrafish nervous system, we disrupted the *grin1a* and *grin1b* genes using CRISPR-Cas9 (Chang *et al.*, 2013; Hwang *et al.*, 2013). We targeted the highly conserved SYTANLAAF motif within the M3 transmembrane segment (Figure 3A). Frameshift mutations in this region will prevent formation of the ion channel as well as the ligand-binding domain (LBD), and likely encode a null allele. We analyzed three germline mutations for each paralogue. Sequencing of each lesion revealed a predicted frameshift mutation and a nearby stop codon (Figures 3B & C). For the majority of our subsequent analysis, we used alleles of *grin1a* and *grin1b* that yielded the earliest predicted stop codon: a 7-nucleotide deletion in *grin1a*^sbu90^ (*grin1a^−/−^*) and a 17-nucleotide insertion in *grin1b*^sbu94^ (*grin1b^−/−^*). RT-PCR confirmed that only *grin1a* or *grin1b* mRNA species containing the indel were present in the respective mutants (Figures 3D & 3E). Given the functional importance of the M3 segment, it is improbable that any cryptic splicing event that bypassed the lesion site in this region could produce a functional receptor. Based on the anticipated consequences of the mutations on NMDAR structure, any protein produced from the *grin1a* and *grin1b* mutant alleles would not form a functional receptor nor conduct current.

**Figure 3.**
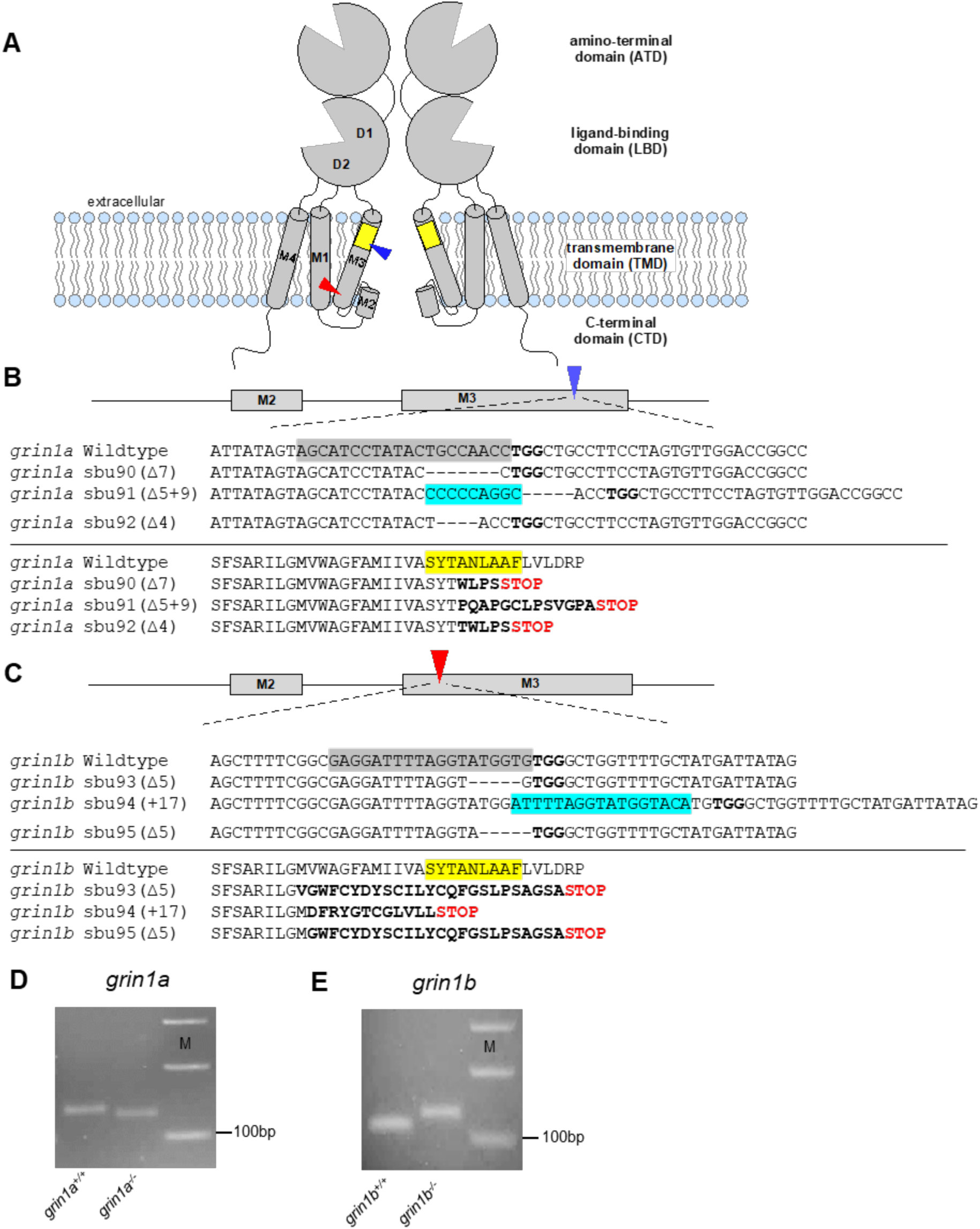
Generation of loss-of-function lesions in *grin1a* and *grin1b* using CRISPR-Cas9. **(A)** Membrane topology of two NMDAR subunits (functional NMDARs are tetramers). Blue and red arrows indicate approximate sites for gRNA targets for *grin1a* and *grin1b*, respectively. NMDARs are composed of four modular domains: the extracellular ATD and LBD; the membrane-embedded TMD; and the intracellular CTD. Each individual subunit contributes three transmembrane segments (M1, M2 & M4) and a M2 pore loop to form the ion channel. The most highly conserved motif in iGluRs, the SYTANLAAF motif (labeled in yellow), is within the M3 segment. gRNAs were designed to prevent generation of this motif and downstream elements (half of LBD, M4 and CTD). **(B & C)** Schematic of gRNA target sites for *grin1a* (blue arrow) **(B)** and *grin1b* (red arrow) **(C)** as well as alignments of nucleotide (*upper*) and amino acid (*lower*) sequences. Induced mutations within the nucleotide sequences are denoted as either dashes (deletions) or highlighted blue (insertions). gRNA target site on the nucleotide sequence (gray highlight) and PAM sites (bolded) are adjacent to generated mutations. Altered amino acid sequence (bolded) and early stop codons (Red STOP) are generated in all alleles, disrupting the SYTANLAAF motif (yellow highlight) as well as removing the D2 lobe of the LBD, which would make the receptor non-functional. **(D & E)** Lesions altered mRNA size in expected fashions. cDNA amplification of: (**D**) *grin1a*^+/+^ and *grin1a* ^−/−^ (unless otherwise noted, *grin1a*^−/−^ denotes the sbu90 allele) producing expected product sizes of 147 bp and 140 bp, respectively; and (**E**) *grin1b*^+/+^ and *grin1b*^−/−^ (unless otherwise noted, *grin1b*^−/−^ denotes the sbu94 allele) producing expected product sizes of 109 bp and 126 bp, respectively. M denotes marker. For all genotypes, RNA was collected from homozygous intercrosses at 3 dpf.

Nonsense mediated decay of mutant mRNA can cause transcriptional adaptation and upregulation of related genes (El-Brolosy *et al.*, 2019). We therefore assessed levels of the lesioned mRNA and the corresponding paralogue in *grin1a*^−/−^ and *grin1b*^−/−^ fish at 3 and 5 dpf (Figure S1). We detected a significant decrease in *grin1a* expression in *grin1a^−/−^* larvae at 3 dpf (Figure S1A), but only a trending decrease at 5 dpf (p = 0.06, Figure S1C), suggesting possible nonsense mediated decay. However, for *grin1b^−/−^* larvae, we detected no corresponding difference in *grin1b* expression (Figure S1B & S1D). To determine whether transcriptional adaptation enhanced expression of the corresponding paralogue, we assayed the expression levels of *grin1a* in *grin1b^−/−^* (Figure S1A & S1C) and of *grin1b* in *grin1a^−/−^* (Figure S1B & S1D) at 3 and 5 dpf, but did not detect any compensation. Hence, transcriptional upregulation of the corresponding paralogue is not a prominent feature of the *grin1* alleles at larval stages.

### *grin1a* and *grin1b* single mutant fish are viable, but only *grin1a^−/−^* display growth defects

Homozygous *grin1a^−/−^* and *grin1b^−/−^* single mutant fish appear morphologically normal at 6 dpf and are recovered at Mendelian ratios (Table S1). Although both survive to adulthood and are fertile, *grin1a^−/−^* adults are not recovered at the expected Mendelian ratios (Table S1). Adult *grin1a^−/−^* fish that are recovered are smaller than their wild-type siblings (Figures 4A & 4C). In contrast, *grin1b^−/−^* fish show no growth deficit (Figures 4B & 4D). For *grin1a^−/−^* fish, a size deficit is apparent between 56 dpf and 98 dpf, though the size difference with their wild-type siblings narrows later in development (Figure 4E). The same reduced viability and growth deficit observed in *grin1a^−/−^* was also present in the *grin1a* sbu92 allele (Table S1 & Figure S2). While we have not determined the cause of increased mortality in *grin1a^−/−^* fish, it may be associated with their slower growth. These phenotypic differences between *grin1a^−/−^* and *grin1b^−/−^* fish indicate that GluN1b is unable to fully compensate for the loss of GluN1a in *grin1a^−/−^*.

**Figure 4.**
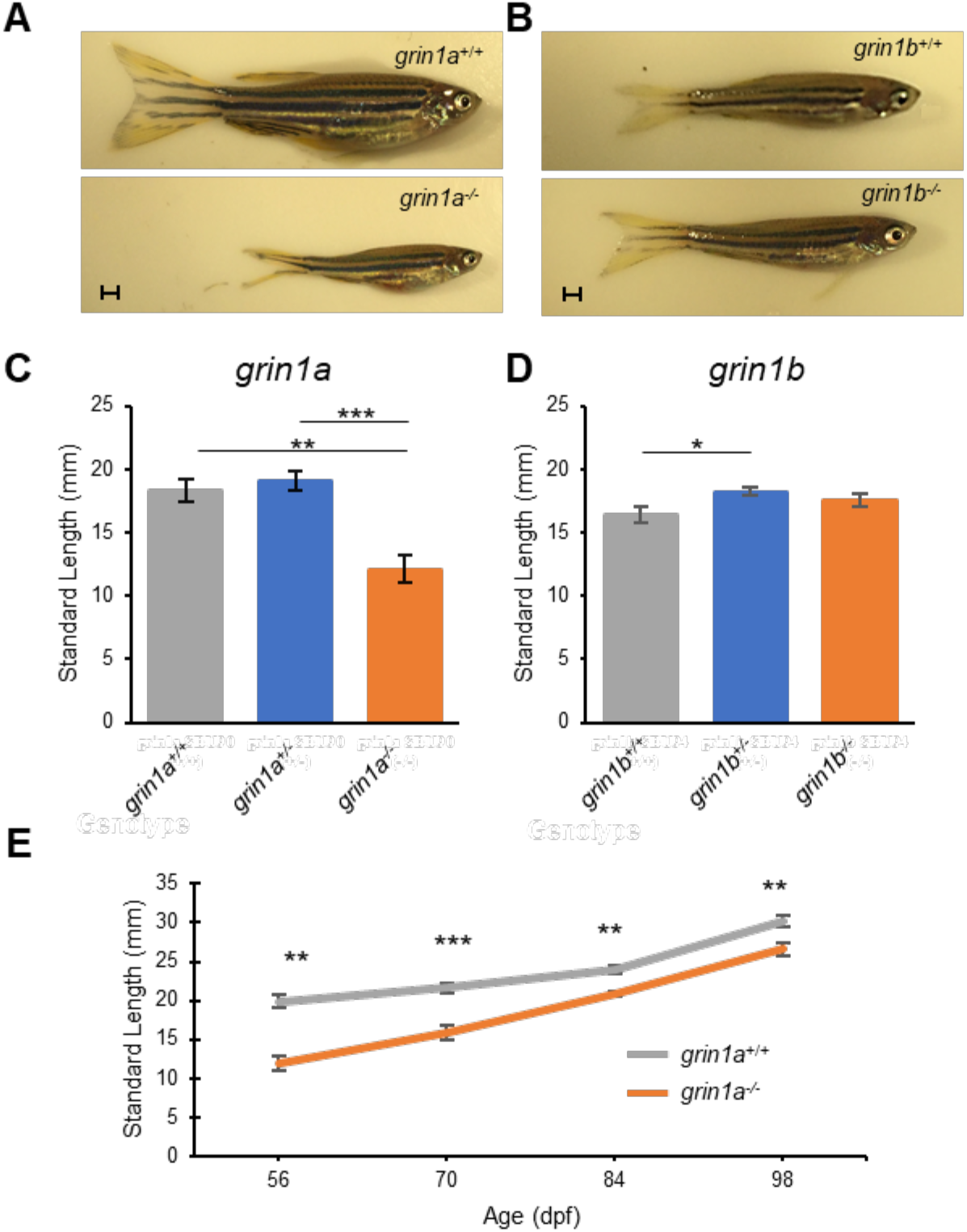
*grin1a*^−/−^, but not *grin1b*^−/−^, fish have decreased growth into adulthood. **(A & B)** Representative photographs of *grin1a*^+/+^ and *grin1a^−/−^* fish **(A)** or of *grin1b*^+/+^ and *grin1b^−/−^* fish **(B)** at 2 months post fertilization. Scale bars, 1 mm. **(C & D)** Standard length comparison (mean ± SEM) of *grin1a*^+/+^ (n = 17), *grin1a*^+/-^ (n = 21), and *grin1a*^−/−^ (n = 5) **(C)** or of *grin1b*^+/+^ (n = 17), *grin1b^+^*^/-^ (n = 62), and *grin1b*^−/−^ (n = 31) **(D)** at 2 months post fertilization. *\*p < 0.05, **p < 0.01, ***p < 0.001, ANOVA, Tukey HSD*. **(E**) Standard length comparison from 8 weeks (56 dpf) to 14 weeks (98 dpf) post fertilization for *grin1a^+^*^/+^ (n = 17) and *grin1a^−^*^/-^ (n = 5). After initial measurement at 8 weeks, fish were moved to individual tanks. \*\**p < 0.01, ***p < 0.001, t-test*.

### Zebrafish lacking *grin1* survive until 10 dpf

Knockout of *grin1* in mice is embryonic lethal (Forrest *et al.*, 1994). Both *grin1a* and *grin1b* single mutant fish survive to adulthood and are reproductively viable. Zebrafish *grin1* double mutants (*grin1a^−/−^; grin1b^−/−^*) are morphologically normal through embryonic and early larval development. The majority of *grin1* double mutants fail to develop a swim bladder (Figure 5A) and we have not been able to recover any past 10 dpf (Table S2). To begin to examine the integrity of the nervous system of the *grin1* double mutants, we used RNA *in-situ* hybridization of a proneural gene (*zash1b/ascl1b)* to assess whether there were gross expressional differences in the *grin1* double mutant neural progenitor population at 3 dpf (Lo *et al.*, 2002). Compared to both wildtype (Figure 5B, left panel) and experimental (Figure 5B*, middle panel*) controls, the regional expression of *zash1b* in the *grin1* double mutants appeared largely unaltered (Figure 5B, right panel). This result suggests that development of zash1b expressing neural progenitors is unaffected by the loss of NMDARs.

**Figure 5.**
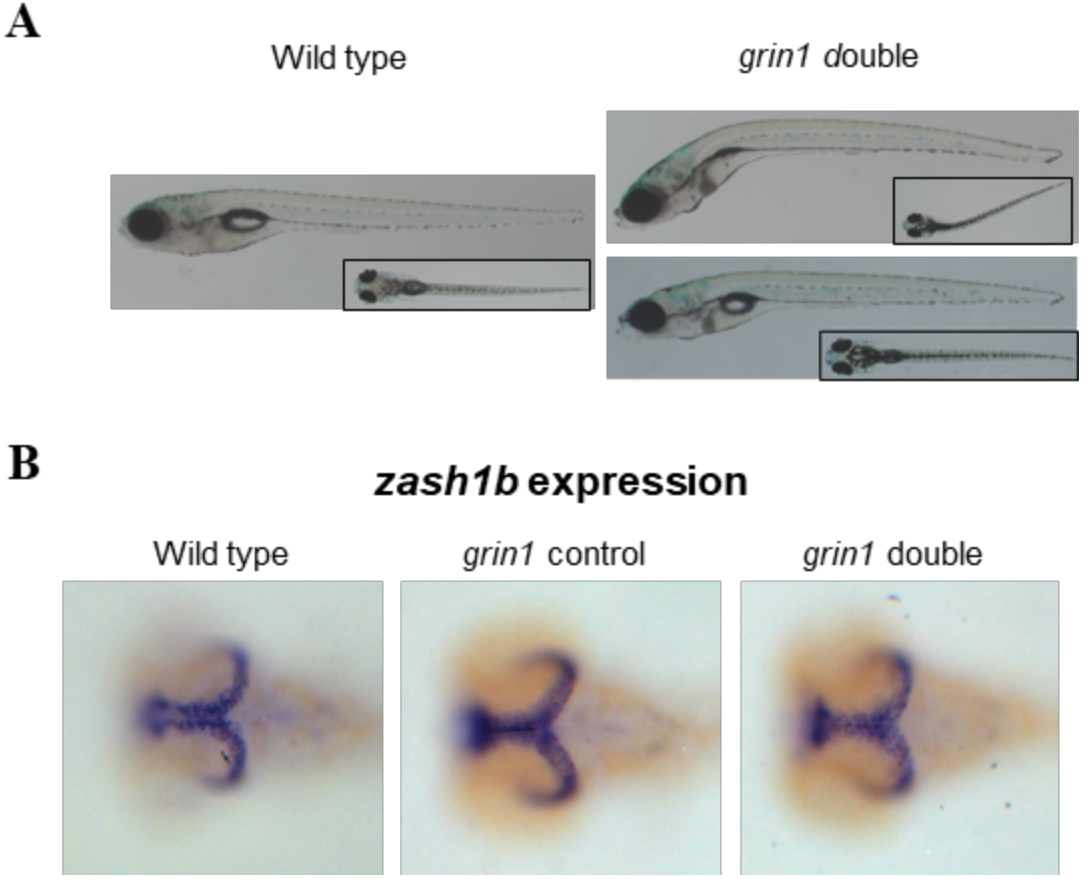
*grin1* double mutants can survive through early larvae development. **(A)** Lateral views of live images of wild type and *grin1a^−/−^*; *grin1b^−/−^* (*grin1* double mutants) fish at 9 dpf. *grin1* double mutants often do not form a swim bladder. **(B)** RNA *in situ* hybridization of *zash1b* for wild type *(left panel)*, *grin1* control (tube control for *in situ; grin1a^+/+^*; *grin1b^−/−^) (middle panel)*, and *grin1* double mutant *(right panel)* at 3 dpf.

### Spontaneous and evoked movements are disrupted in *grin1* double mutant fish but not in single mutants

Zebrafish larvae exhibit a wide repertoire of robust behaviors. The *grin1* double mutant fish provide a unique opportunity to assess the requirement for NMDAR-mediated transmission in generating these behaviors.

Initially, we tested the impact of the loss of NMDARs on spontaneous and evoked swimming movements (Figure 6), because these behaviors provide an overview of basic motor control and response to visual stimuli. We monitored spontaneous movements of individual larva at 6 dpf in 24-well plates during a 50-minute paradigm with periods in light (acclimation and light periods) and dark. Turning off the illumination elicits evoked behavior referred to as a visual motor response (VMR) (Emran *et al.*, 2008). In this paradigm, wild-type zebrafish display stereotyped behaviors (Figures 6A-6D, gray traces): their number of movements stabilizes during the acclimation period and then transiently increases following the transition to darkness. *grin1a^−/−^* (Figures 6A & 6E) and *grin1b ^−/−^* (Figures 6B & 6F) fish replicate this behavior. In contrast, *grin1* double mutant larvae, while they can achieve coordinated swimming, display a different behavioral profile (Figure 6C): they are hyperactive during the initial acclimation period, have reduced levels of activity during the light period, and show a decrease in activity during the dark period (Figures 6C & 6G). Thus, fish lacking both paralogues, and hence void of functional NMDARs throughout development, show an altered phenotype that does not appear in either of the *grin1* single mutants.

**Figure 6.**
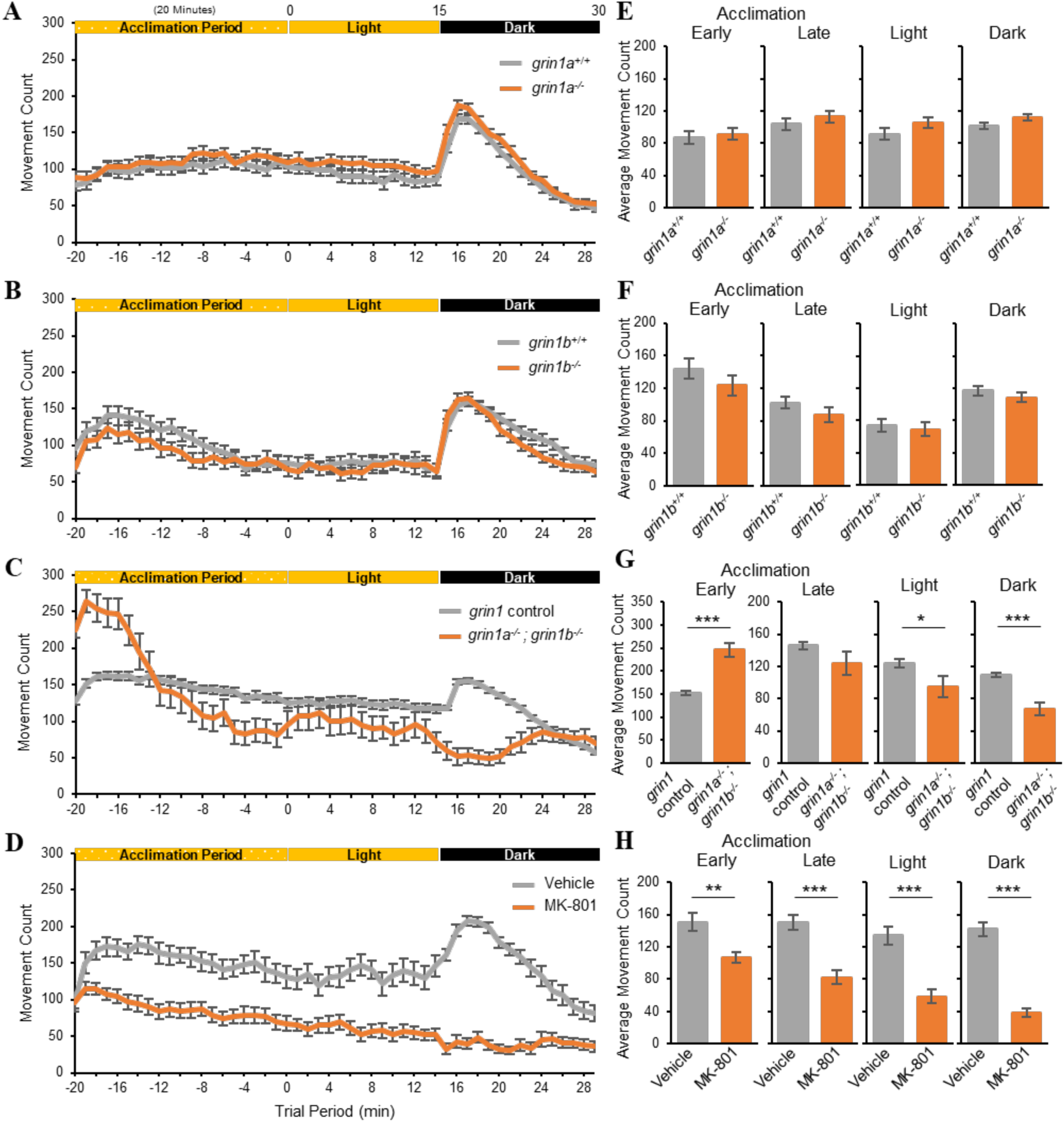
The *grin1* double mutant and MK-801 treated, but not *grin1a^−/−^* or *grin1b^−/−^*, alter spontaneous movement. **(A-D)** Spontaneous movements at 6 dpf and evoked movements after a transition from light to dark. Line graphs of average number of movement counts per minute (mean ± SEM) for: **(A)** *grin1a^+/+^* (n = 64) or *grin1a^−/−^* (n = 79) fish; **(B)** *grin1b^+/+^* (n = 49) or *grin1b^−/−^* (n = 50) fish; **(C)** *grin1* control (*grin1a^+/+^*; *grin1b^+/+^*, *grin1a^+/+^*; *grin1b^+/-^* and *grin1a^+/-^*; *grin1b^+/+^* (n = 148)) or *grin1a^−/−^*; *grin1b^−/−^* (n = 32); and (**D**) fish treated with vehicle (0.1% DMSO) (n = 26) or the NMDAR antagonist MK-801 (20 μM in 0.1% DMSO) (n = 27). Vehicle or MK-801 were acutely administered 1 hour before the start of the spontaneous movement trial and were included in the bath solution throughout the trial. **(E-H)** Bar graphs, of associated spontaneous movement trials, depicting average movements (mean ± SEM) during the early acclimation (first 5 minutes of acclimation), late acclimation (last 15 minutes of acclimation), and light and dark periods for: **(E)** *grin1a^+/+^* or *grin1a^−/−^*; **(F)** *grin1b^+/+^* or *grin1b^−/−^*; **(G)** *grin1* control or *grin1* double mutant; and **(H)** vehicle or MK-801 treated fish. *p < 0.05, **p < 0.01, ***p < 0.001, t-test.

This altered phenotype in *grin1* double mutant fish could reflect that a behavior lacks NMDAR-mediated transmission or that an underlying circuit failed to develop properly due to this lack of transmission. To distinguish these possibilities, we acutely applied MK-801, an NMDAR channel blocker (Traynelis *et al.*, 2010), to wild-type fish (Figure 6D). Because MK-801 treatment occurred only one hour prior to behavioral testing, it would only antagonize NMDAR-mediated transmission and would not alter circuit development. Any phenotype present in the *grin1* double mutants that is recapitulated in acute MK-801 treatment would suggest that the alteration is generated by the lack of NMDAR-mediated transmission, whereas if MK-801 treatment does not replicate the phenotype, it would suggest a developmental effect of the long-term absence of NMDAR-mediated signaling. In the spontaneous and evoked movement paradigm, we observed both outcomes. In parallel with the *grin1* double mutant fish, acute MK-801 treatment significantly reduced spontaneous activity during the light period and removed the stereotypic response to the light change (Figure 6H). In contrast to the *grin1* double mutant fish, acute MK-801 treatment did not induce hyperactivity during the initial acclimation period (Figure 6H).

### MK-801 treatment phenocopies multiple behaviors observed in the *grin1* double mutants

In Figure 6, we measured overall spontaneous and evoked movements and found that in the transition from light-to-dark, *grin1* double mutant fish lacked the stereotypical increase in behavior found in wild-type (Figure 6G), an outcome phenocopied by MK-801 treatment (Figure 6H). Nevertheless, how robust this overlap in behavior is between *grin1* double mutant and MK-801-treated fish is unclear given the gross nature of our movement measurements. We therefore characterized behavior in more detail including the evoked visual motor response in the transition from light-to-dark as well as additional swim parameters during the dark period (Figure 7).

**Figure 7.**
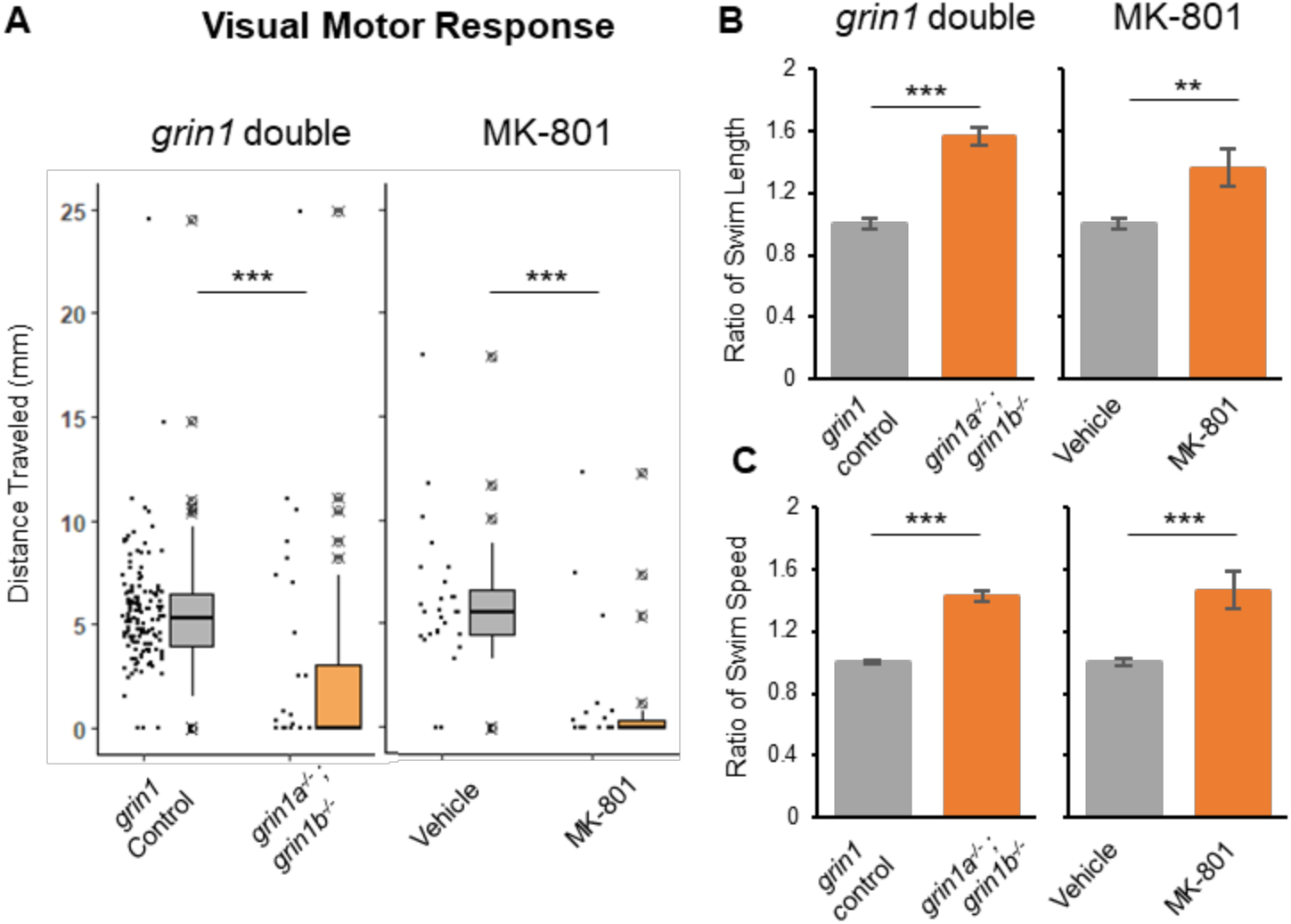
*grin1* double mutants display comparable behaviors in multiple swim parameters to MK-801 treated larvae. **(A)** Box plot and individual responses (dots) for visual motor response (average distance traveled in response to the light change) for *grin1* control (*grin1a^+/+^*; *grin1b^+/+^*, *grin1a^+/+^*; *grin1b^+/-^* & *grin1a^+/-^*; *grin1b^+/+^* (n = 148)) or *grin1* double mutant (*grin1a^−/−^*; *grin1b^−/−^*) (n = 32) (*left panel*); and vehicle (0.1% DMSO) (n = 26) or MK-801 (20 μM in 0.1% DMSO) (n = 27) treated (*right panel)* **(B-C)** Bar graphs (mean ± SEM) for **(B)** Swim length (ratio of distance per movement to control); and **(C)** Swim speed (ratio of speed during movements to control) during the spontaneous locomotion assay at 6 dpf. For calculating swim length and swim speed, only large movements (speeds greater than 8 mm/sec) were used, in order to exclude drifting movement between bursts. Same groups as from panel A. *\*\*p < 0.01, ***p < 0.001, t-test*.

To assay the visual motor response, we measured the distance traveled (mm) in the second after the change from light-to-dark. In response to this light change, *grin1* double mutant fish and MK-801 treated fish showed significant decreases in their visual motor response compared to controls (Figure 7A). The majority of these fish had a completely extinguished VMR, with a few responding outliers. *grin1a* and *grin1b* single mutants showed no alteration in VMR (Figure S3A). These results parallel the findings of gross motor response to the light change (Figures 6C & 6D).

To characterize additional swim parameters that were not visually evoked, we measured the swim length (distance (mm) per movement) (Figure 7B) and swim speed (distance (mm) per time (s) of movements) (Figure 7C) during the dark period. We found that both the swim length and swim speed were significantly increased in both the *grin1* double mutant and the MK-801 treated fish compared to controls. *grin1a* and *grin1b* single mutants showed no alteration in these swim kinetics (Figures S3B & S3C). Thus, on the transition from light-to-dark, MK-801 completely phenocopies the effects of the grin1 double mutation, suggesting at minimum these deficits reflect a lack of NMDAR-mediated transmission.

### Behavioral defects in the *grin1* double mutants with a developmental origin

As observed in the spontaneous locomotion paradigm, the *grin1* double mutant fish (Figures 6C & 6G), but not MK-801 treated fish (Figure 6D & 6H), exhibited hyperactivity immediately after being placed in the testing apparatus. We assume this hyperactivity reflects NMDAR-mediated disruption in brain development and is not dependent on NMDAR-mediated transmission per se. If this hyperactivity is of developmental origin, we would anticipate to see it at earlier developmental periods. Indeed, *grin1* double mutant fish at earlier developmental time points (4 and 5 dpf) also show hyperactivity during the early acclimation period (Figures S4A & S4B). To test if this hyperactivity was dependent on NMDAR-mediated activity and whether it reflected any off-target effects of MK-801, we applied acute MK-801 treatment to the *grin1* double mutant fish, but found that MK-801 did not abolish the hyperactivity in the early acclimation period (Figure S5). These findings are consistent with the idea that a circuit underlying the early acclimation period is disrupted due to the lack of NMDAR-mediated signaling during development.

Zebrafish larvae are thigmotaxic, spending less time in the center of the well compared to the periphery (Figure 8A) (Schnorr *et al.*, 2012). We therefore tested if *grin1* double mutant fish showed a similar behavior. While *grin1* double mutants enter the center of the well as often as their control siblings (Figure 8B, left panel), they exit more rapidly and do not remain in the center as long as controls (Figure 8C, left panel). Fish treated with MK-801, however, show no difference in either their entrances into the center of the well, or the time spent there, as compared to controls (Figures 8B & 8C, right panels). These observations expand upon the scope of behaviors with a potential developmental origin.

**Figure 8.**
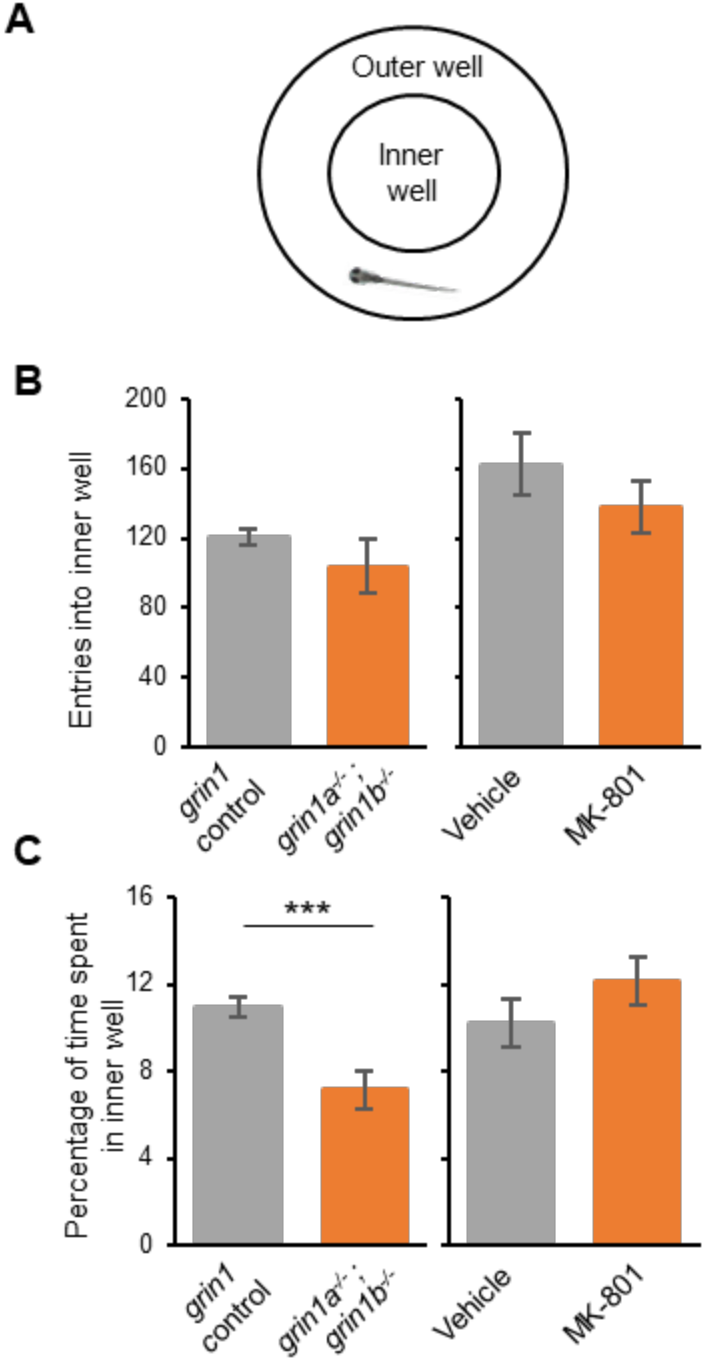
MK-801 treatment cannot phenocopy all altered behaviors in *grin1* double mutants. **(A)** Schematic representation of the inner and outer portions of a behavior well **(B & C)** Number of entries into the inner section of behavior well (mean ± SEM) **(B)** and percentage of time spent in the inner behavior well (mean ± SEM) **(C)** during the spontaneous movement paradigm (Figure 6). Different conditions/genotypes tested include: *grin1* control (*grin1a^+/+^*; *grin1b^+/+^*, *grin1a^+/+^*; *grin1b^+/-^* and *grin1a^+/-^*; *grin1b^+/+^* (n = 148) or *grin1a^−/−^*; *grin1b^−/−^* (n = 32); and vehicle (0.1% DMSO) (n = 26) or MK-801 (20 μM in 0.1% DMSO) (n = 27) treated.

### NMDARs are required for normal feeding behavior

Prey capture is a complex behavior requiring visual and motor coordination (Semmelhack *et al.*, 2014). To determine the involvement of NMDARs in this behavior, we used a larval feeding assay to test the ability of individual larvae to capture paramecia (Gahtan *et al.*, 2005) (see Material & Methods). Because of variation in the number of paramecia eaten by control larvae from day to day, we normalized the proportion of paramecia eaten by the experimental group to their same-day control (Figure 9).

**Figure 9.**
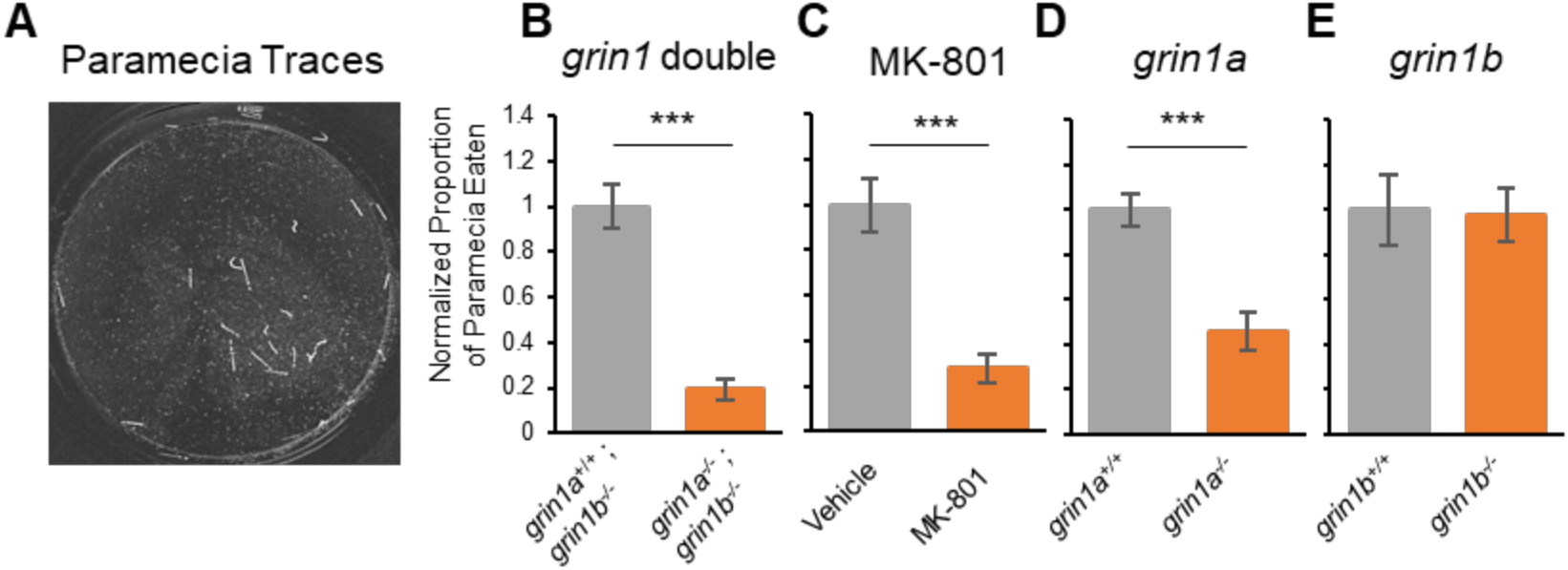
*grin1* double mutants and MK-801 treatment lead to feeding deficits. **(A)** Representative traces of paramecium movement used to assay feeding. Traces generated by analyzing 2.5 seconds of paramecia movement. **(B -E)** Proportion of paramecia eaten eaten over the trial period (mean ± SEM) normalized to control for**: (B)** *grin1a*^+/+^; *grin1b*^−/−^ (n = 18) or *grin1a^−^*^/-^; *grin1b*^−/−^ (n = 33) fish; **(C)** fish treated with vehicle (n = 15) or MK-801 (n = 15); **(D)** *grin1a*^+/+^ (n = 25) or *grin1a*^−/−^ (n = 23) fish; and **(E)** *grin1b*^+/+^ (n = 14) or *grin1b*^−/−^ (n = 10) fish. For vehicle and MK-801 treated, larvae were batch exposed to either vehicle (0.1% DMSO) or MK-801 (20 μM in 0.1% DMSO) for 2 hours prior to start of assay. *\*\*\*p < 0.001, t-test*.

Compared to the controls, *grin1* double mutant fish showed a significant deficit in prey capture (Figure 9B). It is noteworthy that *grin1* double mutants, presumably lacking all NMDAR-transmission, can sufficiently coordinate their sensory and locomotor activity to successfully hunt paramecium (Movie 1), albeit in a diminished capacity. These deficits in prey capture could be replicated by acute exposure of wild-type larvae to MK-801, likely indicating a requirement for NMDAR-mediated transmission in generating this behavior (Figure 9C).

Contrary to all previously tested behaviors, decreases in prey capture were also observed in *grin1a^−/−^* larvae (Figure 9D). To support this surprising finding, we also observed this feeding deficit in the *grin1a* sbu92 allele (Figure S6). No deficits were observed in *grin1b^−/−^* larvae (Figure 9E). These findings indicate that there are differences in the roles of the two zebrafish GluN1 paralogues in generating more complex behaviors.

### *grin1* double mutants fail to habituate to acoustic stimuli

NMDAR signaling is associated with nonassociative learning behaviors in larval zebrafish (Wolman *et al.*, 2011). Habituation to acoustic stimuli in larval zebrafish is robust. In order to assess habituation, we first determined their short latency startle (SLC) responsiveness (Burgess & Granato, 2007). SLC responsiveness was increased in the *grin1* double mutants compared to controls (Figure 10A). Paralleling this result, SLC responsiveness was increased in *grin1a^−/−^* larvae compared to heterozygous and wild-type sibling controls at low and high intensity stimuli (Figure 10B). In contrast, *grin1b^−/−^* larvae showed no differences in SLC responsiveness (Figure 10C).

**Figure 10.**
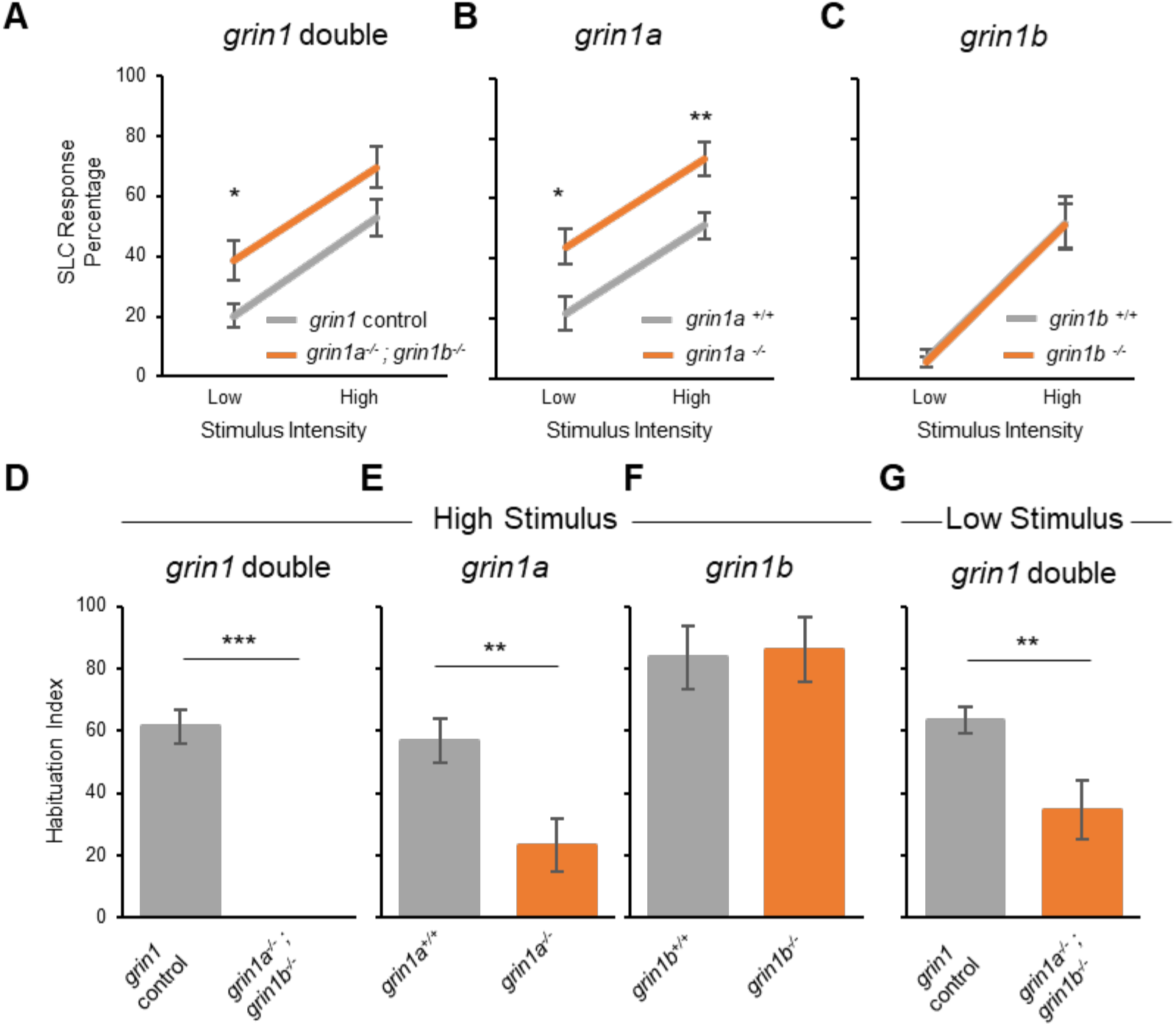
Disruption of sensorimotor gating in *grin1* mutant fish. **(A-C)** Responsiveness of larvae to an acoustic startle of low and high frequency measured as percentage of larvae responding with a short latency c-bend (SLC) for: **(A)** *grin1* control (n = 22) or *grin1* double mutant (n = 20); **(B)** *grin1a*^+/+^ (n = 18) or *grin1a^−^*^/-^ (n = 20); and **(C)** *grin1b*^+/+^ (n = 13) or *grin1b^−^*^/-^ (n = 15). *\* = p < 0.05, ** = p < 0.01, t-test.* **(D-F)** Habituation to a high intensity acoustic stimuli measured, as a habituation index over the course of the trial for: **(D)** *grin1* control (n = 12) or *grin1* double mutant (n = 4); **(E)** *grin1a*^+/+^ (n = 22) or *grin1a^−^*^/-^ (n = 21); and **(F)** *grin1b*^+/+^ (n = 12) or *grin1b^−^*^/-^ (n = 10). *\* = p < 0.05, ** = p < 0.01, t-test*. **(G)** Habituation to a low intensity acoustic stimuli, measured as a habituation index over the course of the trial for *grin1* control (n = 42) or *grin1* double (n = 14).

We then tested *grin1a* and *grin1b* single mutants for changes in habituation. We presented startling stimuli at 1s intervals and measured habituation as the percent decrease in SLC responsiveness. Wild-type siblings exhibited 57 ± 7% (mean ± SEM) habituation, *grin1a^−/−^* larvae showed only 23 ± 8% (Figure 10E), whereas *grin1b^−/−^* larvae had no deficits in habituation (Figure 10F). To test for additive effects of *grin1a* and *grin1b*, we then tested *grin1* double mutants for changes in habituation. Double mutants showed a decrease in habituation compared to *grin1a^+/+^;grin1b*^−/−^ and *grin1a^+/-^;grin1b*^−/−^ controls (Figure 10D). This decrease was not significantly different from *grin1a* single mutants (independent samples t-test, t(31) = 0.9, p = 0.11). These findings are consistent with previous work that has shown that MK-801 treatment leads to both increased response to acoustic startle and decreased habituation (Wolman *et al.*, 2011). Together, these data indicate that in zebrafish GluN1 (primarily GluN1a) is required for acoustic startle responses and non-associative learning.

## DISCUSSION

We have developed a zebrafish model to interrogate NMDAR function throughout early development. NMDAR-mediated signaling is integral to generating many behaviors and higher order brain functions including neural plasticity, learning and memory, and neurodevelopment (Paoletti *et al.*, 2013). However, connections between developmental functions of NMDARs and subsequent impacts on behaviors have been limited, at least in part, due to the lethality of mouse models prior to stages when behavior can be studied (Forrest *et al.*, 1994). To explore this phase of NMDAR function, we studied the obligatory GluN1 subunit during early zebrafish development.

We investigated the expression and function of the two *grin1* paralogues in zebrafish, *grin1a* and *grin1b*, that arose from an ancient genome duplication (Amores *et al.*, 1998). Heterologous expression of zebrafish GluN1 paralogues in HEK293 cells demonstrated that in terms of key NMDAR properties, current conductance and Ca^2+^ permeability, zebrafish GluN1a and GluN1b are similar to each other and to the rat GluN1 subunit (Figure 2). Although the ion channel function of GluN1a and GluN1b were largely similar, the expression domains of zebrafish *grin1a* and *grin1b* highlighted potentially important differences (Figure 1). *grin1a* is expressed more robustly at 1 dpf than *grin1b* and it is expressed in the developing spinal cord where *grin1b* is absent, suggesting differential spatiotemporal requirements for the two paralogues.

We find that fish lacking all NMDAR mediated transmission (*grin1* double mutants) survive until 10 dpf. These fish displayed abnormal spontaneous and evoked behavior (Figure 6) including reduced visual motor responses (Figure 7A), prey capture deficits (Figure 9) and a failure to habituate to acoustic stimuli (Figure 10). Still, the *grin1* double mutant fish executed a variety of synchronized behaviors including burst swimming behavior, responses to light and acoustic stimuli, and the ability to capture prey though at diminished rates.

It is unlikely that any residual GluN1 function remains in the *grin1* double mutants based on several lines of evidence. First, we were unable to detect any wild-type *grin1* mRNA in the mutant (Figure 3D). Given the location of the lesions, any alternative splicing event that circumvented the mutations would bypass the highly conserved M3 pore lining domain, which would render the channel non-functional. Lastly, many features of the *grin1* double mutant can be phenocopied with the NMDAR antagonist MK-801 (Figures 7 & 8) and MK-801 does not alter behavior phenotypes of the *grin1* double mutant (Figure S5). Based on these results, and the well-established requirement for GluN1 in NMDAR-mediated neurotransmission, we conclude that the *grin1* double mutants lack all NMDAR function.

Compensatory mechanisms may act in the GluN1 mutants to restore some functions. Genetic compensation occurs in response to nonsense mediated decay (El-Brolosy *et al.*, 2019), but we did not detect any compensatory increases in the other paralogue (Figure S1). While NMDARs have distinct functional properties, other classes of iGluRs, including AMPA and kainate receptors, could provide some compensatory function either at the synaptic or circuit level. Future studies to explore compensatory mechanisms in the *grin1* mutants are needed to account for the finding that *grin1* double mutant fish can execute complex behaviors (e.g., prey-capture).

While many *grin1* double mutant phenotypes are replicated by short-term administration of MK-801, which suggests acute effects of loss of NMDAR-mediated transmission, others are not, for example, hyperactivity upon first entering the apparatus (Figure 6C) and a reduced time spent in the center of the well (Figure 8). These phenotypes may stem from developmental effects of loss of NMDAR-mediated transmission. In addition, both of these phenotypes may have a common origin, being result of anxiogenic stimuli. Nevertheless, more detailed investigations are needed to fully address this issue.

Acute MK-801 treatment phenocopies the prey-capture deficits of the *grin1* double mutant (Figure 9), which indicates that NMDAR-mediated transmission is at least required to generate this behavior. However, prey capture is a complex behavior and the short term block in NDMAR mediated transmission may also be overshadowing additional developmental roles of NMDAR. Loss of only *grin1a* is sufficient to reduce the prey capture ability. This finding may begin to explain both the reduction in viability, and the growth deficits observed in *grin1a^−/−^* adults (Figure 4 & Table S1). This finding also suggests that *grin1a* and *grin1b* have unique functions, as removal of *grin1b* did not alter prey capture ability.

The *grin1* double mutant displayed multiple phenotypes (Figure 10) previously observed in pharmacologically inhibition of NMDAR in larvae: an increase in SLC responsiveness seen with MK-801 treatment (Bergeron *et al.*, 2015), and a decrease in habituation to an acoustic stimuli as seen with both MK-801 and ketamine (Roberts *et al.*, 2011; Wolman *et al.*, 2011). Interestingly, this phenotype appears to be caused solely by loss of *grin1a* (Figure 10). The acoustic startle circuit generation of SLC responses is well characterized and may be an ideal setting to elucidate potential functional or expressional differences between *grin1a* and *grin1b* (Lopez-Schier, 2019).

We have developed a model that provides a unique opportunity to explore the requirements for NMDARs in early vertebrate nervous system. Given the similarity in both sequence and measured physiological properties of the *grin1* paralogues to their mammalian orthologues, zebrafish are an ideal organism to study the developmental roles of NMDAR. We found that GluN1 and by extension NMDAR mediated transmission required for multiple behaviors including the visual motor response, prey capture and habituation to acoustic stimuli. Further studies are needed to further elucidate these, and other, developmental roles of GluN1, as well as understanding potential compensatory mechanisms allowing for coordinated movements in these larval zebrafish.

