## Supplemental Material for "A model to study NMDA receptors in early nervous system development"

**Zoodsma et al**

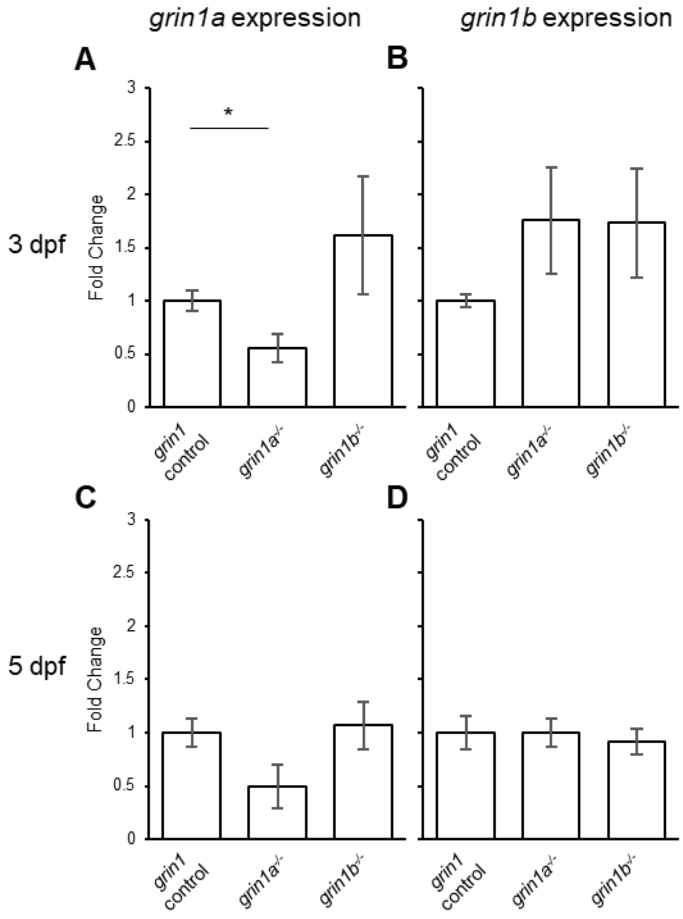

**Supplemental Figure 1. Limited nonsense mediated decay and no compensation of the paralogous *grin1* mRNA at early developmental timepoints (relates to Figure 3).**

**(A & B)** Quantitative PCR (qPCR) of *grin1a* **(A)** and *grin1b* **(B)** expression at 3 dpf in *grin1* control (n = 9), *grin1a*$^{-/-}$ (n = 9) or *grin1b*$^{-/-}$ (n = 9) fish. None of the values were significantly different in (B).

**(C & D)** Quantitative PCR (qPCR) of *grin1a* **(C)** and *grin1b* **(D)** expression at 5 dpf in *grin1* control (n = 6), *grin1a*$^{-/-}$ (n = 6) or *grin1b*$^{-/-}$ (n = 6) fish. None of the values were significantly different in either (C) or (D).

**p < 0.05, t-test.*

To assess potential nonsense mediated decay, we assayed the level of *grin1a* expression in *grin1a*$^{-/-}$ fish (Figures S1A & S1C, *left two bars*) and *grin1b* expression in *grin1b*$^{-/-}$ fish (Figures S1B & S1D, *rightmost bar compared to the control, leftmost bar*). We observed a significant decrease of *grin1a* expression in *grin1a*$^{-/-}$ fish at 3 dpf (Figure S1A), suggesting possible nonsense mediated decay. However, we did not observe a significant decrease at 5 dpf (Figure S1C), nor did we observe any such decrease in *grin1b*$^{-/-}$ fish (Figure S1B & S1D).

To assess potential compensation, we assayed the levels of *grin1a* expression in *grin1b*$^{-/-}$ fish (Figures S1A & S1C, *rightmost bar compared to the control, leftmost bar*) and of *grin1b* expression in *grin1a*$^{-/-}$ fish (Figures S1C & S1D, *rightmost bar compared to the control, leftmost bar*). At these early developmental time points, we did not observe compensation of the paralogous gene at the level of mRNA.

| Allele | Age Screened | Number Screened | Expected Percentage | Percentage Wildtype | Percentage Mutant | Mutant to Wildtype Ratio | chi–squared P-value |
|---|---|---|---|---|---|---|---|
| *grin1a* sbu90 | 6 dpf | 1240 | 25.0 % | 24.0 % | 26.1 % | 1.09 : 1 | 0.812 |
| | Adult | 292 | 25.0 % | 30.1% | 11.9 % | 0.39 : 1 | 1.55e-6 *** |
| *grin1a* sbu92 | 6 dpf | 274 | 25.0 % | 26.6 % | 24.1% | 0.90 : 1 | 0.812 |
| | Adult | 159 | 25.0 % | 27.0 % | 8.1 % | 0.33 : 1 | 8.66e-6 *** |
| *grin1b* sbu94 | 6 dpf | 548 | 25.0 % | 24.8 % | 24.5 % | 0.99 : 1 | 0.936 |
| | Adult | 172 | 25.0 % | 20.3 % | 27.3 % | 1.34 : 1 | 0.359 |

**Supplemental Table 1: *grin1a*$^{-/-}$, but not *grin1b*$^{-/-}$, adults have decreased viability (relates to Figure 4).**

Viability of *grin1a* and *grin1b* alleles genotyped at 6 dpf and after 2 months (adults) as recorded from heterozygous intercrosses for each allele. Expected recovery of both wildtype and mutant alleles from heterozygous intercrosses is 1/4. Both the sbu90 and sbu92 alleles for *grin1a* are recovered at a significantly reduced percentage as adults.

****$p < 0.001$, chi squared test.*

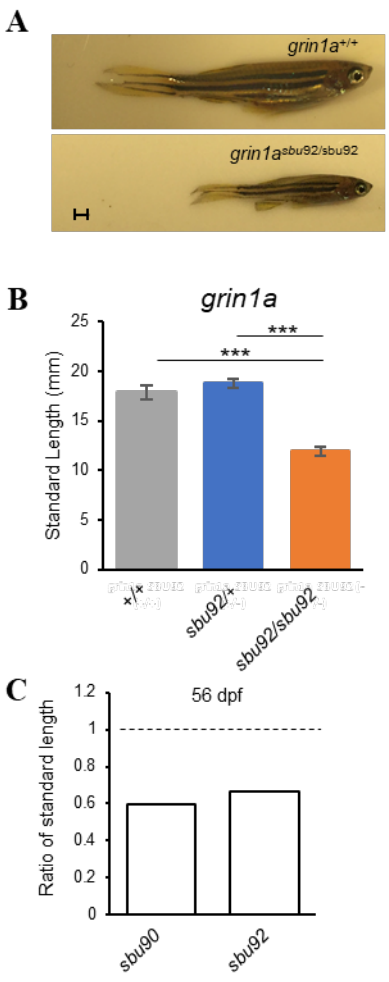

**Supplemental Figure 2. *grin1a* sbu92 allele recapitulates attenuated growth phenotype of *grin1a* sbu90 (relates to Figure 4).**

**(A)** Representative photographs of *grin1a*$^{+/+}$ and *grin1a*$^{sbu92/sbu92}$ fish at 2 months post fertilization.

**(B)** Standard length comparison (mean ± SEM) of *grin1a*$^{+/+}$ (n = 17), *grin1a*$^{sbu92/+}$ (n = 35), and *grin1a*$^{sbu92/sbu92}$ (n = 7) at 2 months post fertilization. ****$p < 0.001$, ANOVA, Tukey HSD.*

**(C)** Ratio of standard length of *grin1a*$^{-/-}$ to *grin1a*$^{+/+}$ at 8 weeks (56 dpf) for *grin1a* sbu90 and sbu92 alleles (Table S1).

The *grin1a* sbu92 allele recapitulates the growth phenotype, a decrease in size at 2 months, observed in *grin1a* sbu90 (Figure 4). The ratio of sbu90 or sbu92 to wildtype size at 2 months is indistinguishable (Figure S2C).

| Allele | Age Screened | Number Screened | Expected Percentage | Percentage Wildtype | Percentage Mutant | Mutant to Wildtype Ratio | chi–squared P-value |
|---|---|---|---|---|---|---|---|
| ***grin1* Double** | 6 dpf | 557 | 6.25 % | 5.2 % | 6.3 % | 1.21 : 1 | 0.094 |
| | Adult | 117 | 6.25 % | 7.7 % | 0.0 % | 0 : 1 | 0.00345 ** |

**Supplemental Table 2. Fish with both *grin1a* and *grin1b* (*grin1* double mutant) eliminated do not survive to adulthood (relates to Figure 5).**

Viability of *grin1 double* genotyped at 6 dpf and after 2 months (adults) was generated from a double heterozygous intercross using the sbu90 and sbu94 alleles for *grin1a* and *grin1b,* respectively. Expected recovery of fish mutant for both alleles for a double heterozygous intercross is 1/16. *grin1* double mutants were recovered in Mendelian ratios at 6 dpf but were unrecoverable as adults.

*$**p < 0.01$, Chi Squared Test.*

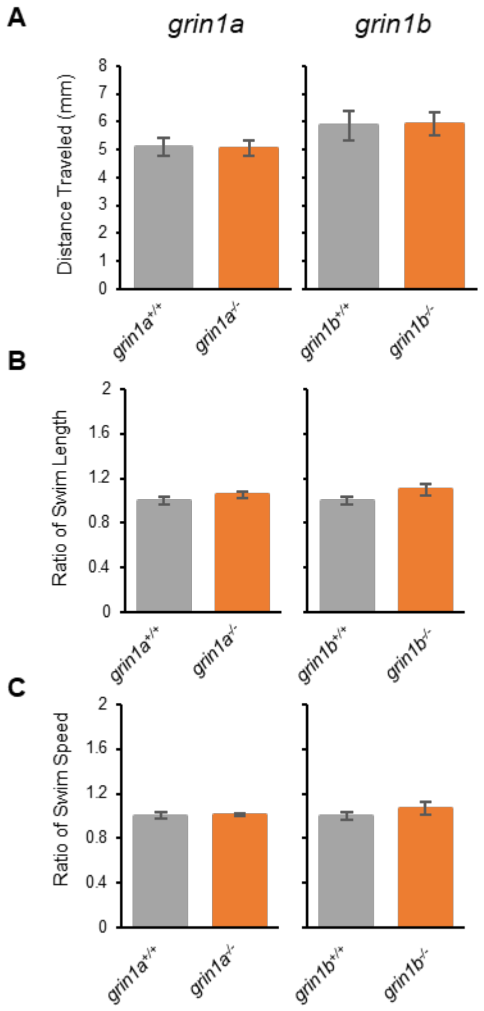

**Supplemental Figure 3. *grin1* single mutants show no alterations in additional swim parameters (relates to Figure 6 & 7).**

**(A-C)** Bar graphs (mean ± SEM) for: **(A)** Visual motor response (average distance traveled in response to the light change); **(B)** Swim length (ratio of distance per movement to control); and **(C)** Swim speed (ratio of speed during movements to control) during the spontaneous locomotion assay at 6 dpf. For calculating swim length and swim speed, only large movements (speeds greater than 8 mm/sec) were used, in order to exclude drifting movement between bursts. Different conditions/genotypes tested include: *grin1a*$^{+/+}$ (n = 64) or *grin1a*$^{-/-}$ (n = 79) *(left panels)*; *grin1b*$^{+/+}$ (n = 49) or *grin1b*$^{-/-}$ (n = 50) (*right panels)*

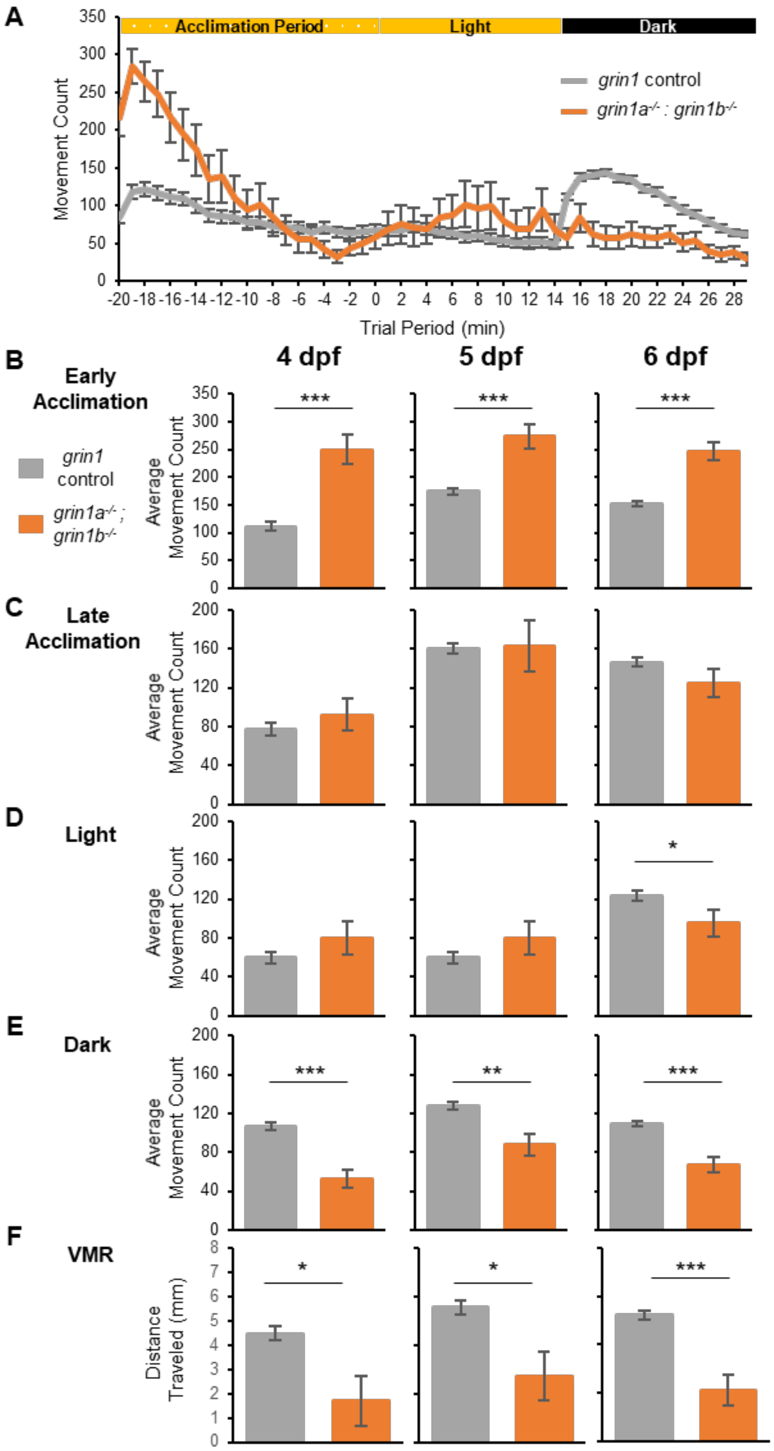

**Supplemental Figure 4. *grin1* double mutants show similar spontaneous movements throughout development (relates to Figures 6 & 7).**

**(A)** Spontaneous movements at 4 dpf and evoked movements after a transition from light to dark. Line graphs of average number of movement counts per minute (mean ± SEM) for *grin1* control (*grin1a*$^{+/+}$; *grin1b*$^{+/+}$, *grin1a*$^{+/+}$; *grin1b*$^{+/-}$, and *grin1a*$^{+/-}$; *grin1b*$^{+/+}$ (n = 101)) or *grin1a*$^{-/-}$; *grin1b*$^{-/-}$ (n = 17).

**(B-F)** Bar graphs depicting average movements (mean ± SEM) during: (**B**) early acclimation (first 5 minutes of acclimation); (**C**) late acclimation (last 15 minutes of acclimation); **(D)** light and **(E)** dark periods; and **(F)** visual motor response (VMR) for 4 dpf (*grin1* control (n = 101) and *grin1a*$^{-/-}$; *grin1b*$^{-/-}$ (n = 17)) (*left panels*); 5 dpf (*grin1* control (n = 100) and *grin1a*$^{-/-}$; *grin1b*$^{-/-}$ (n = 17)) (*center panels*), and 6 dpf (*grin1* control (n=148) and *grin1a*$^{-/-}$; *grin1b*$^{-/-}$ (n = 32) (*right panels*).

*p < 0.05, **p < 0.01, ****p < 0.001, t-test*.

*grin1* double mutant larvae show similar behavioral profiles at 4, 5 and 6 dpf compared to controls. Data for 6 dpf are also shown in Figure 6. As seen at 6 dpf (Figure 6), the hyperactivity in the early acclimation period (Figure S4A & S4B) is also present at these earlier developmental time points. Gross movement response to the light change (Figure S4E) and visual motor response (Figure S4F) are also conserved at these developmental time points.

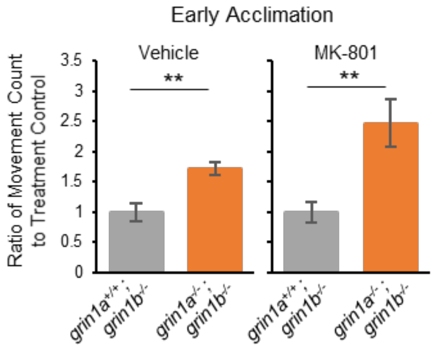

**Supplemental Figure 5. MK-801 treatment does not abolish hyperactivity during the early acclimation period in *grin1* double mutant fish (relates to Figure 6 & 8).**

Ratio of average movement count (mean ± SEM) during early acclimation normalized to treatment control for *grin1a*$^{+/+}$; grin1b$^{-/-}$ (n = 13 & 14) and *grin1a*$^{-/-}$; *grin1b*$^{-/-}$ (n = 8 & 9) for vehicle (0.1% DMSO) or MK-801 (20 µM in 0.1% DMSO) treatments respectively.

*** = $p < 0.01$, t-test.*

*grin1* double mutant fish show hyperactivity during the early acclimation period (Figure 6C), which we interpret to reflect a developmental change. Consistent with this idea is that an acute application of MK-801 does not block this hyperactivity.

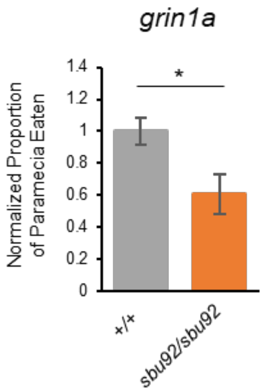

**Supplemental Figure 6. *grin1a* sbu92 allele recapitulates the prey capture deficit seen in sbu90 (relates to Figure 9).**
Proportion of paramecium eaten over the trial period (mean ± SEM) normalized to control for *grin1a*$^{+/+}$ (n = 13), *grin1a*$^{sbu92/+}$ (n = 18), and *grin1a*$^{sbu92/sbu92}$ (n = 11) fish. Data were collected as in Figure 9. *$*p < 0.05$, t-test.*

Both the *grin1a* sbu90 (Figure 9) and sbu92 alleles show a prey capture deficit in the paramecia feeding behavioral paradigm.
